# Loss of the systemic vitamin A transporter RBPR2 affects the quantitative balance between chromophore and opsins in visual pigment synthesis

**DOI:** 10.1101/2024.07.08.602543

**Authors:** Rakesh Radhakrishnan, Matthias Leung, Anjelynt Lor, Swati More, Glenn P. Lobo

## Abstract

The distribution of dietary vitamin A/all-*trans* retinol (ROL) throughout the body is critical for maintaining retinoid function in peripheral tissues and for generating visual pigments for photoreceptor cell function. ROL circulates in the blood bound to the retinol binding protein 4 (RBP4) as RBP4-ROL. Two membrane receptors, RBPR2 in the liver and STRA6 in the eye are proposed to bind circulatory RBP4 and this mechanism is critical for internalizing ROL into cells. Here, we present a longitudinal investigation towards the importance of RBPR2 and influence of the diet on systemic retinoid homeostasis for visual function. Age matched *Rbpr2*-KO (*Rbpr2^-/-^*) and wild-type (WT) mice were fed either a vitamin A sufficient (VAS) or a vitamin A deficient (VAD) diet. At 3- and 6-months, we performed retinoid quantification of ocular and non-ocular tissues using HPLC analysis and complemented the data with visual physiology, rhodopsin quantification by spectrophotometry, and biochemical analysis. At 3-months and compared to WT mice, *Rbpr2^-/-^* mice fed either vitamin A diets displayed lower scotopic and photopic electroretinogram (ERG) responses, which correlated with HPLC analysis that revealed *Rbpr2^-/-^* mice had significantly lower hepatic and ocular retinoid content. Interestingly, with the exception of the liver, long-term feeding of *Rbpr2^-/-^* mice with a VAS diet promoted all- *trans* retinol accumulation in most peripheral tissues. However, even under VAS dietary conditions significant amounts of unliganded opsins in rods, together with decreased visual responses were evident in aged mice lacking RBPR2, when compared to WT mice. Together, our analyses characterize the molecular events underlying nutritional blindness in a novel mouse model and indicate that loss of the liver specific RBP4-ROL receptor, RBPR2, influences systemic retinoid homeostasis and rhodopsin synthesis, which causes profound visual function defects under severe vitamin A deficiency conditions.

## INTRODUCTION

Vitamin A/all-*trans* retinol (ROL) has pleiotropic functions in the human body, attributable to its several biologically active forms^1^. These processes include vision, corneal development, immune system functioning, maintaining epithelium integrity, cellular growth and differentiation, fetus and central nervous system development^1–5^. Dietary vitamin A is the precursor for the visual chromophore (11-*cis* retinaldehyde/11-*cis* retinal) and all-*trans* retinoic acid (a*t*RA). The vitamin A active metabolite in the eye 11-*cis* retinal binds with photoreceptor opsin, a G-coupled protein receptor, to form rhodopsin that is a critical pigment for light perception^4,5^. Upon light exposure, 11-*cis* retinal is isomerized to all-*trans* retinal, causing a photobleaching process where rhodopsin forms several different intermediate states that trigger a G-protein signaling pathway. The vitamin A metabolite a*t*RA is a hormone-like molecule that regulates gene expression through interactions with nuclear receptors that are critical for the differentiation and patterning of the eyes. Vitamin A deficiency in the eye leads to impaired night vision due to deficient rhodopsin formation and if left untreated can cause photoreceptor cell death and blindness^4^. Thus, an understanding of mechanisms that facilitate and regulate the uptake, transport, and long-term storage of dietary vitamin A/ROL for systemic retinoid homeostasis is significant to the design of strategies aimed at attenuating retinal degenerative diseases associated with ocular ROL deficiency in conditions like Retinitis Pigmentosa or Leber Congenital Amaurosis^6–18^.

The main transport form of dietary vitamin A in the circulation to peripheral tissues is all-*trans* retinol (ROL), which is bound to the retinol binding protein 4 (RBP4) as holo-RBP4 (RBP4-ROL)^1,2,19–20^. Two membrane receptors that bind to RBP4 and facilitate the internalization of ROL into tissues from the circulation have been proposed. Previously, biochemical and genetic evidence have shown the involvement of Stimulated by Retinoic Acid 6 (STRA6) that is highly expressed in the retinal pigmented epithelium (RPE), in the uptake of circulatory RBP4-ROL into the eye^21–23^. STRA6 is however not expressed in major peripheral/non-ocular tissues, including the liver that functions as the main storage organ for dietary vitamin A^2,24–29^. This indicates that alternate vitamin A receptors might exist in such tissues, which may be responsible for whole-body retinoid homeostasis in the support of chromophore production. First identified by the Graham laboratory in 2013, the retinol binding protein 4 receptor 2, RBPR2 or STRA6like (STRA6l), is implicated in the systemic tissue uptake and storage of ROL from RBP4^30^. Our previous biochemical and genetic analysis of RBPR2 in cell lines, and in zebrafish and mice deficient in RBPR2, showed that it was a bona fide RBP4-vitamin A receptor and involved in ROL internalization^26–28^. Additionally, we have previously showed that RBPR2 contains specific RBP4-ROL binding motifs and loss of the systemic vitamin A transporter, RBPR2, in mice resulted in visual function defects^24,25^. However, it is yet unknown what long-term effects of the diet are in *Rbpr2^-/-^* mice in maintaining systemic and ocular retinoid homeostasis, and on the quantitative relationship between chromophore and opsins in the generation of rhodopsin for visual function and in maintaining retinal health.

To answer these questions, we have now conducted a longitudinal study in *Rbpr2^-/-^* and WT mice fed with either a vitamin A sufficient (VAS) or vitamin A deficient (VAD) diet, to better understand the consequences of the diet and genetics on systemic vitamin A uptake, whole-body retinoid and ocular retinoid homeostasis. We also aimed to investigate the long-term effects of the diet under these genotypes on chromophore production for visual function. By comparing RBPR2-deficient mice to control mice under different conditions of dietary vitamin A supply, we studied the effects of the systemic RBP4-ROL receptor, RBPR2, on visual pigment biogenesis and photoreceptor cell function. Even though RBPR2 is not expressed in the eye, we observed that mice lacking the systemic vitamin A receptor, RBPR2, display characteristic features of not only systemic retinoid deficiency, but also lower ocular retinoid levels, which result in decreased phototransduction. We further show that in the absence of RBPR2 and under long-term dietary vitamin A restriction, these mice are more susceptible to ocular consequences of vitamin A deprivation, which included lower rhodopsin concentrations, delayed kinetics in rod and cone opsin regeneration under scotopic and photopic conditions, and the presence of unliganded opsins in rod photoreceptors, altogether affecting visual function.

Together, our study demonstrates the importance of the systemic RBP4-ROL receptor, RBPR2, for maintaining liver retinoid homeostasis that is important for the critical balance between chromophore and opsins in visual pigment synthesis. Our study establishes the RBP4-ROL transporter, RBPR2, as an important component of whole-body retinoid homeostasis and mammalian visual function, especially under fasting or restricted dietary vitamin A conditions, where it plays a critical role in preventing retinal pathologies associated with ocular vitamin A deprivation.

## MATERIALS AND METHODS

### Materials

All chemicals unless stated otherwise were purchased from Sigma-Aldrich (St. Louis, MO, USA) and were of molecular or cell culture grade quality.

### Animals, animal husbandry, and diets

*Rbpr2*-knockout (*Rbpr2^-/-^*) mice used in the study have been previously described^24^. Six-week-old wild type (WT) mice (C57BL/6J) were purchased from JAX labs. Breeding pairs and litters of *Rbpr2^-/-^* and WT mice were genotyped and found to be negative for the known *Rd8* and *Rd1* mutations, as previously described by us^24,31^. Breeding pairs of mice were fed purified chow diets containing 8 IU of vitamin A/g (Research diets) and provided water ad libitum and maintained at 24°C in a 12:12 hour light-dark cycle. All animal experiments were approved by the Institutional Animal Care and Use Committee (IACUC) of the University of Minnesota (protocol # 2312-41637A), and performed in compliance with the ARVO Statement for the use of Animals in Ophthalmic and Vision Research. Post weaning (P21), equal numbers of male and female mice were randomly distributed to vitamin A feeding groups. For experiments, WT or *Rbpr2^-/-^* mice (n=16 per group) were fed purified rodent diets (AIN-93G; Research Diets, New Brunswick, NJ) containing the recommended 4 IU of vitamin A/g (Vitamin A sufficient diet; VAS) or specially formulated and purified low vitamin A /vitamin A deficient (VAD) diets contained 0.22 IU vitamin A/g based on the AIN-93G diet (Research Diets, New Brunswick, NJ)^10–12,24,34^ for up to 6-months. The percentage difference of dietary vitamin A between the two diets is ∼180%.

### Immunohistochemistry and Fluorescence Imaging

Mice at 3- and 6-months on vitamin A diet conditions were euthanized by CO2 asphyxiation and cervical dislocation. Eyes were enucleated and fixed with either 4% PFA in 1X PBS or in Davidson’s fixative for 4 hours at 4^0^C. Paraffin-embedded retinal sections (∼8 µm) were processed for antigen retrieval and immunofluorescence. Primary antibodies were diluted in 1% normal goat serum (NGS) blocking solution as follows: anti-rhodopsin 1D4 for mouse rod opsin (1:200; Abcam, St. Louis, MO, USA), R/G cone opsins (1:100; Millipore, St. Louis, MO, USA), and 4′,6-diamidino-2-phenylendole (DAPI; 1:5000, Invitrogen) or Hoechst (1:10,000, Invitrogen) to label nuclei. All secondary antibodies (Alexa Fluor 488) were used at 1:5000 concentrations (Molecular Probes, Eugene, OR, USA). In addition, the TrueVIEW vector autofluorescence quenching kit SP-8500-15 was used to help reduce background noises and non-specific stains. Optical sections were obtained with a Nikon AXR confocal and processed with the Nikon Viewer software, or using a Keyence BZ-X800 scope. All fluorescently labeled retinal sections on slides were analyzed by the BioQuant NOVA Prime Software (R & M Biometrics, Nashville, TN, USA) and fluorescence within individual retinal layers were quantified using Image J or Fiji (NIH). After mounting, images were captured using 40X and 60X objectives. The acquired retinal images were calibrated with the ZEISS ZEN 3.4 software package and intensities were quantified and data were plotted in GraphPad Prism.

### Electroretinogram (ERG) Analysis

#### Dark-adapted Scotopic ERG

Mice were dark-adapted for 12-16 hours. Under single source red light, eyes were dilated with tropicamide and phenylephrine before being sedated with Isoflurane using a calibrated Isoflurane machine (equipped with precision vaporizer, flow meter, and oxygen). A drop of 2.5 percent Hypromellose ophthalmic demulcent solution (Systane) was placed onto the cornea just before the electroretinogram. Dark adapted electroretinograms (ERG) were obtained (0.01 to 100 cd s/m2) using a Celeris instrument (Diagnosys). Before running a combined dark and light protocol (Diagnosys Espion software), the electrodes were placed on the cornea and an impedance was measured. The Rod response recovery after bleaching, Celeris ERG protocols were performed with pulse frequency 1 and pulse intensity 1. The protocol was set to acquire prebleach amplitudes and to initiate the bleaching protocol using Light adaptation time 180secs. After bleaching the amplitudes of the *a*- and *b*-waves were measured under scotopic conditions (1 cd s/m2) for every minute till 10 minute post bleaching. The curves were plotted in GraphPad Prism and the half-life were measures using the formula Y=(Y0-NS)*exp(-K*X) + NS. K is the rate constant in inverse units of the X axis. The half-life equals the ln(2) divided by K.

#### Light-adapted Photopic ERG

The light adapted mice were sedated and eyes were dilated and the ERG electrodes were placed as mentioned in dark adapted mice methods section. To measure the photoreceptor Cone response *a*, *b*-wave and retinal ganglion response photopic Negative Response ERG, Celeris ERG protocol phNR were performed under light adapted condition with pulse frequency 2 and pulse intensity 20[P] and background intensity 40[P] and color green. The amplitudes were recorded and plotted in GraphPad Prism.

### Purification of Rhodopsin and absorbance spectroscopy

Mice were dark-adapted for 12-16 hours. Under single source red light, the mice were then (CO2) euthanized, and the retina was harvested surgically. In strict dark conditions, the retinal tissues were homogenized with 20 mM bis-tris propane, 150mM NaCl, and 1mM EDTA buffer pH 7.5 with protease inhibitor. The homogenates were centrifuged 15min at 16000g refrigerated. The supernatants were discarded, and the pellets solubilized for 1 hour on a rotating platform at 4^0^C in 20 mM bis-tris propane, 150mM NaCl, 20 mM n-dodecyl-b-D-maltoside (DDM) buffer pH 7.5 with protease inhibitor. The lysate was centrifuged for 1 hour at 16000g at 4^0^C. The supernatant was incubated for 1 hour with 30 µL HighSpec Rho1D4 MagBeads (Cat 33299 Cube Biotech Germany). The resin was washed on a magnetic stand with 20 mM bis-tris propane, 500 NaCl pH7.5 buffer two times and three times with low salt 20 mM BTP, 100 NaCl, and 2 mM DDM buffer. The VAPA peptides dissolved in low salt buffer were used at a concentration of 0.1mg/mL volume 60 µL to elute the Opsins from 1D4 resin. The eluted Opsin was analyzed on an Agilent Cary 60 spectrophotometer Instrument; the measured absorbances were plotted in GraphPad Prism Version 10.1 and calculated for free Opsin using a 280 nm/500 nm absorbance ratio^32^, using the extinction coefficient ε_500_ = 40,600 M^−1^ cm^−1^. The concentration of ligand-free opsin was calculated using the extinction coefficient ε_280_ = 81,200 M^−1^ cm^−1^.

### High-Performance Liquid Chromatography (HPLC) analyses of retinoids

Retinoid isolation procedures was performed under a dim red safety light (600 nm) in a dark room. Animals were first euthanized with CO_2_ asphyxiation, and pertinent tissues were removed from the carcass. The tissue was the homogenized in 0.9% saline with a handheld tissue grinder, consisting of a glass tube and glass pestle. Methanol (2 mL) was added into the homogenate to precipitate the proteins within the homogenate. The retinoid content from the tissue homogenate was then extracted with 10 mL of hexane (twice), with the aqueous layer subsequently removed. The combined hexane extracts were then evaporated with a vacuum evaporator, resuspended in 100 μL of hexane, then injected into an HPLC for analysis. HPLC analysis was performed on an Agilent 1260 Infinity HPLC with a UV detector. The HPLC conditions employed two normal-phase Zorbax Sil (5 μm, 4.6 × 150 mm) columns (Agilent, Santa Clara, CA, USA), connected in series within the Multicolumn Thermostat compartment. Chromatographic separation was achieved by isocratic flow of mobile phase containing 1.4% 1-Octanol/2% 1,4-Dioxane/11.2% Ethyl Acetate/85.4% Hexane, at a flow rate of 1 ml/min for 40 minutes. Retinaldehydes, Retinols, and Retinyl esters were detected at 325 nm using a UV-Vis DAD detector, while the UV absorbance spectra was collected from 200 nm – 700nm. For quantifying molar amounts of retinoids, the HPLC was previously calibrated with synthesized standard compounds and as previously described by us^24^. Calculation of concentration (µM): Standards were injected in concentrations ranging from 0-3.5µM prepared solutions in the mobile phase. The plotted concentrations were fit through linear regression to obtain R-equation (y=mx+c) where y is the peak area (mAU*sec); m is the slope of the calibration curve and c is the y-intercept. The area from the HPLC peaks of the samples (mAU*sec) are interpolated into concentration and expressed as picomoles. For eyes the values are expressed as picomoles/eye; for Liver the values are expressed as picomoles/mg; For Serum the values are expressed as picomoles/microliter.

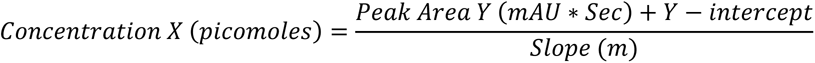

### Statistical Analysis

Data were expressed as means ± standard error mean, statistical analysis by ANOVA and student *t*-test. Differences between means were assessed by Tukey’s honestly significant difference (HSD) test. P-values below 0.05 (p<0.05) were considered statistically significant. For western blot analysis, relative intensities of each band were quantified (densitometry) using the Image *J* software version 1.54 and normalized to the loading control β-actin. The qRT-PCR analysis was normalized to 18S RNA, and the ΔΔCt method was employed to calculate fold changes. Data of qRT-PCR were expressed as mean ± standard error of mean (SEM). Statistical analysis was carried out using GraphPad Prism v 10.1.

## RESULTS

### Design of mouse studies and dietary vitamin A intervention

In this study, we used the previously established RBPR2-deficient (*Rbpr2^-/-^*) mouse line and isogenic C57BL/6J wild-type (WT) mice^24^. Breeding pairs and litters of *Rbpr2^-/-^* and WT mice were genotyped and found to be negative for the known *Rd8* and *Rd1* mutations. Breeding pairs were fed a breeder chow diet containing 8 IU vitamin A/g, to avoid developmental complications. Each group had males and females and were of the same C57BL/6J background. Groups of WT and *Rbpr2^-/-^* mice were fed either a vitamin A sufficient (VAS; 4 IU vitamin A/g) or vitamin A deficient (VAD; 0.22 IU/g) diet post weaning at P21, which are custom diets routinely used to control vitamin A status in mice^34,35^. The percentage difference of dietary vitamin A between the two diets is ∼180%. Additionally, the 4 IU retinol/g concentration in the VAS diet is consistent with the recommended vitamin A intake for rodents and corresponds to 1.2 mg retinol activity equivalent (RAE), which is also a recommended intake in humans. After 3-months of dietary intervention, the first cohort of mice were sacrificed to determine the short-term effects, while the second cohort of mice were sacrificed after 6-months of dietary intervention to determine the long-term effects of vitamin A deficiency on systemic all-*trans* retinol uptake and liver storage and on ocular health in the different genotypes (**Figure 1**).

**Figure 1:**
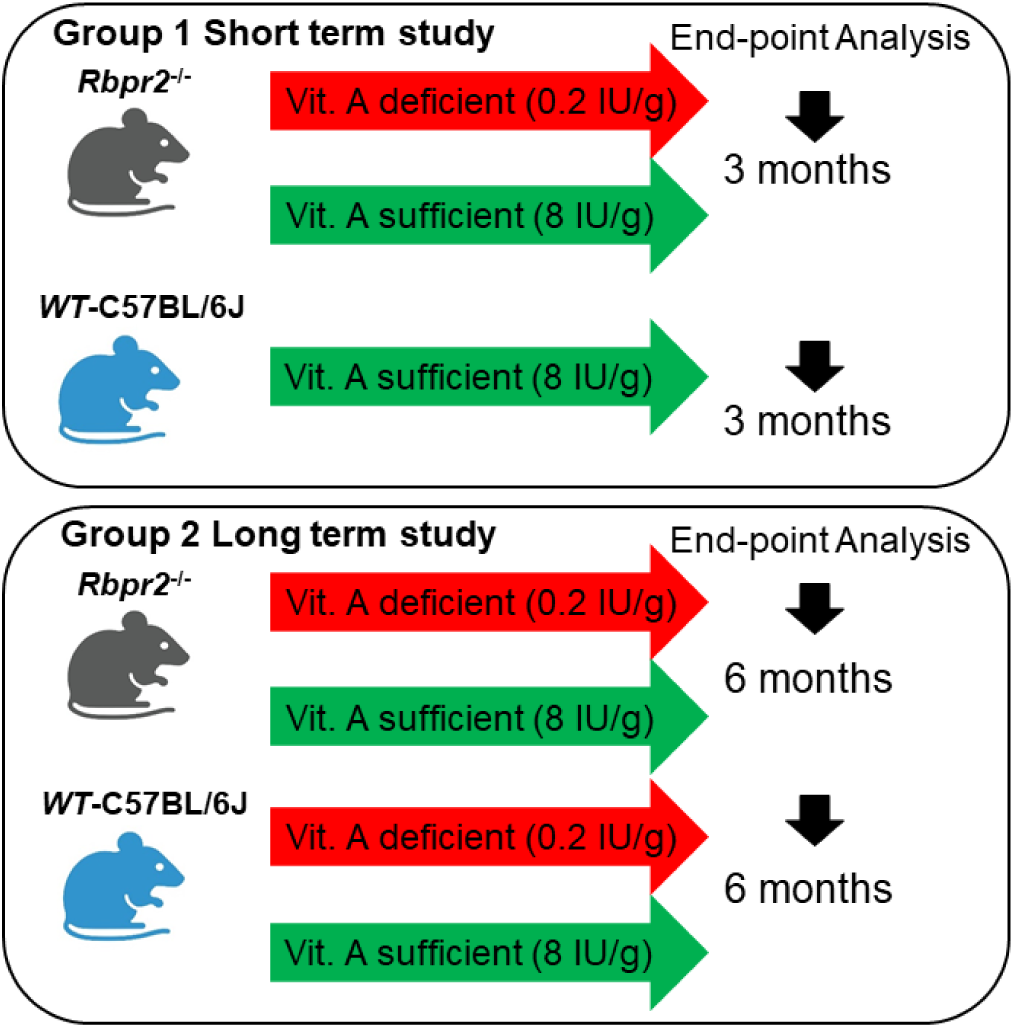
Schematic overview of the mouse study and vitamin A diet intervention. At P21, we fed cohorts of WT and *Rbpr2^-/-^* mice with a vitamin A sufficient (VAS) or vitamin A deficient (VAD) diet. At the 3-month (early time point analysis) and 6-month time point (late time point analysis), non-invasive tests for retinal function and integrity were performed. Ocular and multiple non-ocular tissues were harvested and subjected to High performance liquid chromatography (HPLC) analysis to quantify all-*trans* ROL and retinoid concentrations.

### Effect of the diet on systemic and ocular all-*trans* retinol levels in *Rbpr2^-/-^* mice

Unlike STRA6, the vitamin A receptor, RBPR2, is highly expressed in the liver and to a lesser extent in systemic/ non-ocular tissues to support dietary vitamin A storage and re- absorption of circulatory ROL from RBP4-ROL^2,20,24,25,30,36,37,38^. Thus, the RBPR2 receptor is proposed to regulate whole-body retinoid homeostasis, which is important to the supply of all-*trans* ROL to the eye for maintaining ocular rhodopsin levels for phototransduction, especially under fasting conditions^2,30,37^. We first examined how global loss of RBPR2 in mice affects systemic all-*trans* retinol (ROL) levels in ocular and non-ocular tissues by high-performance liquid chromatography (HPLC) analysis and compared these results to age-matched wild-type (WT) mice. At the early time-point, in 3-month old *Rbpr2^-/-^* mice on either VAS or VAD diets, all-*trans* ROL levels in the liver and eye were significantly lower (p<0.005 and p<0.05 respectively), when compared to age-matched WT mice on VAS diets (**Figures 2A, 2B, 2E, 2F, Supplemental Figures S1 and S2**). Additionally, total retinoids in liver and eyes of *Rbpr2^-/-^* mice on VAD diets, were lower to those observed in *Rbpr2^-/-^* or WT mice on VAS diets (**Figures 2C and 2G, Supplemental Figures S1 and S2**). Similarly, when we measured all-*trans* ROL levels in various non-ocular tissues we observed lower ROL concentrations in *Rbpr2^-/-^* mice on VAD diets, compared to WT mice on VAS diets (**Supplemental Figure S3**). These results indicate that in the absence of RBPR2 and under vitamin A restriction, *Rbpr2^-/-^* mice are more susceptible to vitamin A deficiency, which affects liver and eye retinoid stores (**Figure 3**).

**Figure 2:**
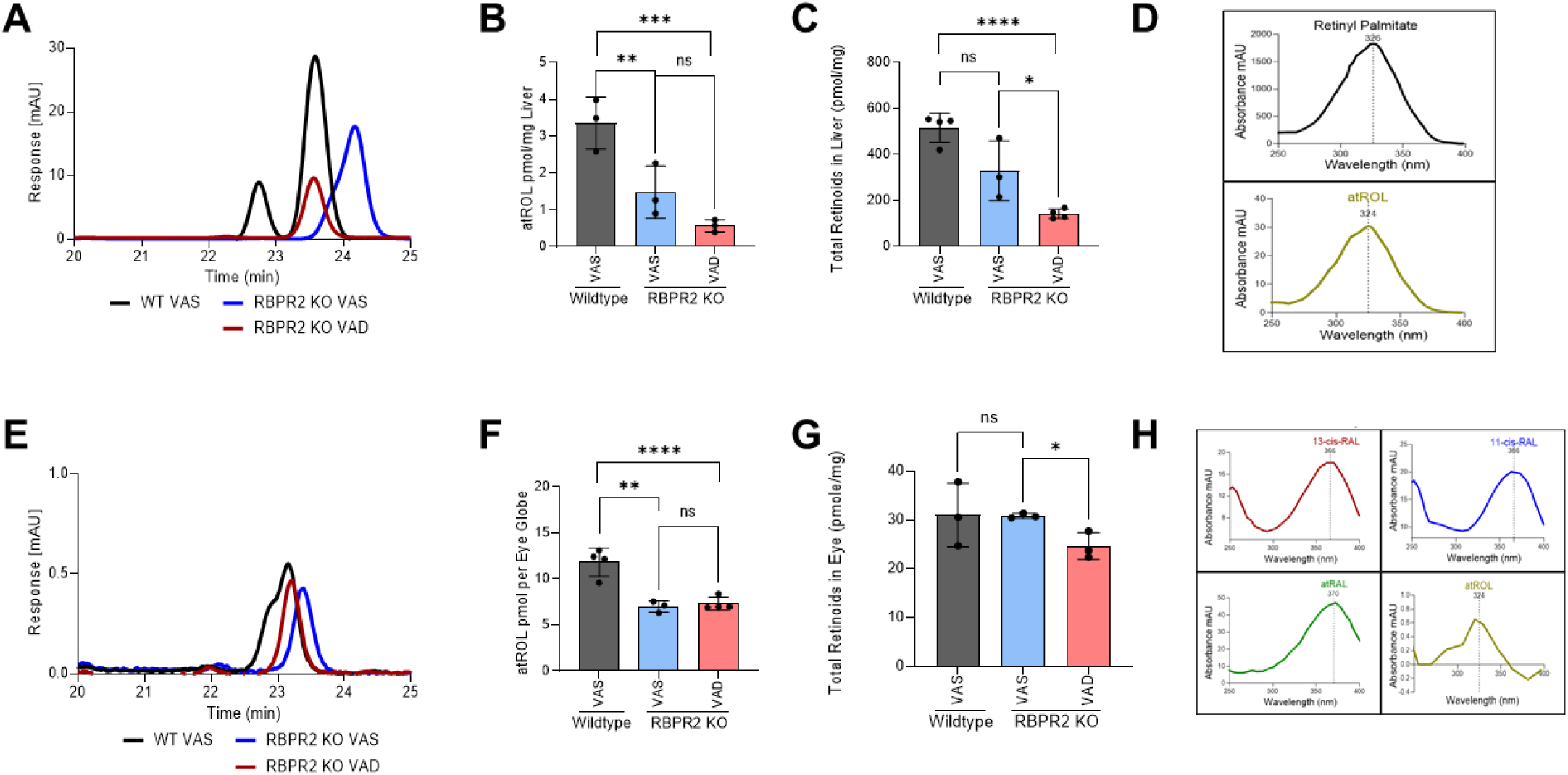
Quantification of all-*trans* ROL and total retinoids in mice tissue at 3- months of age. High performance liquid chromatography (HPLC) analysis and quantification of all-*trans* ROL and total retinoid content in the liver (**A-C**), and eyes (**E-G**), isolated from WT and *Rbpr2^-/-^* mice at 3-months of age on VAS or VAD diets. Absorbance peaks of individual retinoids in the liver (**D**) and eye (**H**). Values are presented as ±SD. Student *t*-test, *p<0.05; **p<0.005; ***p<0.001; ****p<0.0001. VAS, vitamin A sufficient diet; VAD, vitamin A deficient diet; WT, wild-type mice. n=3-4 animals per group.

**Figure 3:**
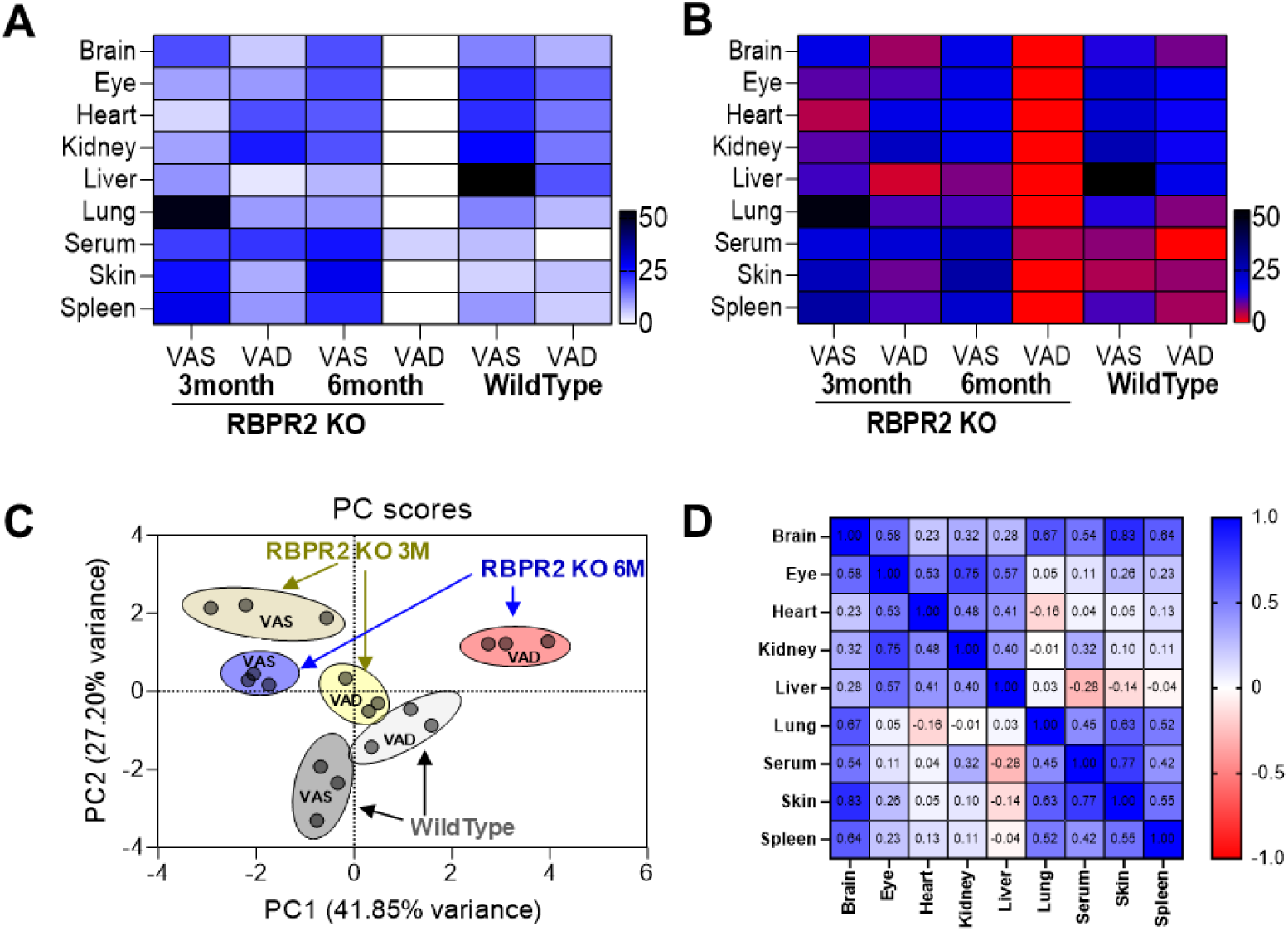
All-*trans* ROL distribution in ocular and non-ocular tissues of WT and *Rbpr2^-/-^* mice on VAS or VAD diets. (**A**) Heat map of all-*trans* ROL levels in WT and *Rbpr2^-/-^* mice at 3- and 6-months of age, in various tissues. (**B, C**) data clustering and variance by Principal Component Analysis (PCA). (**D**) Correlation matrix showing the all-*trans* ROL distribution pattern in tissue samples. VAS, vitamin A sufficient diet; VAD, vitamin A deficient diet; WT, wild-type mice.

We then investigated the long-term effects of the diet in *Rbpr2^-/-^* and WT mice on ROL levels in the liver and eye. In WT mice under either diets, ROL levels in the liver were not significantly affected, however, total retinoids in liver were lower in WT mice under VAD diets, indicating that stored retinoids were likely being distributed under vitamin A restriction (**Figures 4A-C**). In *Rbpr2^-/-^* mice under either diets, ROL and total retinoid concentrations in liver were lower to those of WT mice (**Figures 4A-C**). The total mass loss of ROL stores in *Rbpr2^-/-^* mice under either diet was ∼80% lower compared to WT mice under VAS diets (**Figure 4D**). In the eye of WT mice at the 6-month time point, while ROL levels were lower under VAD conditions, total ocular retinoids concentrations were unchanged. Similarly, 11-*cis* retinal levels in the eye in WT mice under VAD conditions were higher than WT mice under VAS diets, indicating that ROL was being converted to 11-*cis* RAL under vitamin A deficiency in WT mice, which also corresponds to lower ocular ROL levels in these mice under VAD diets (**Figures 4F and 4G**). Conversely, in *Rbpr2^-/-^* mice under VAD conditions, ocular ROL and total retinoid levels were significantly lower to WT mice under VAS or VAD or *Rbpr2^-/-^* mice under VAS conditions (**Figures 4E-G**). Additionally, 11-*cis* retinal levels in *Rbpr2^-/-^* mice under VAD diets, were significantly lower those WT mice under VAD or VAS conditions (**Figure 4G**). Similarly, when we measured all-*trans* ROL levels in various non-ocular tissues at the 6- month time point we observed lower ROL concentrations in these tissues of *Rbpr2^-/-^* mice on VAD diets, compared to *Rbpr2^-/-^* mice on VAS diet or WT mice on VAD or VAS diets (**Supplemental Figure S4**). These results suggest a major role for RBPR2 in maintaining ROL and total retinoid homeostasis in the liver and other non-ocular/ systemic tissues for RBP4-ROL uptake and re-distribution to the eye especially under fasting conditions (**Figure 3B**).

**Figure 4:**
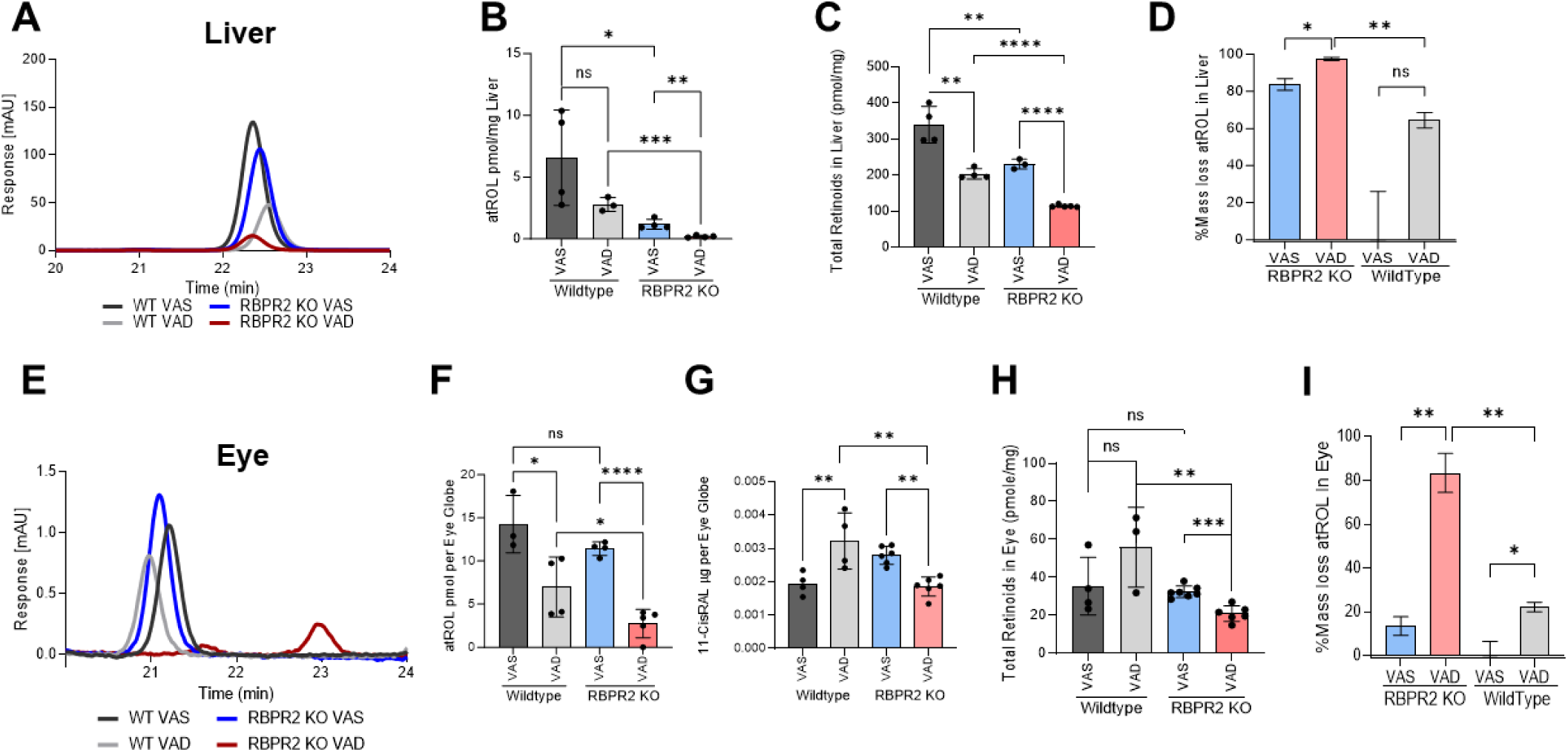
Quantification of all-*trans* ROL and total retinoids in mice tissue at 6- months of age. High Performance Liquid Chromatography (HPLC) was used to determine the all-*trans* ROL and total retinoid content in the liver (**A-C**) and eyes (**E-G**) isolated from WT and *Rbpr2^-/-^* mice at 6-months of age, which were fed either a VAS or VAD diet. Quantification of percentage mass loss of all-*trans* ROL in liver (**D**) and eyes (**H**) of *Rbpr2^-/-^* vs. WT mice. Values are presented as ±SD. Student *t*-test, *p<0.05; **p<0.005; ***p<0.001; ****p<0.0001. VAS, vitamin A sufficient diet; VAD, vitamin A deficient diet; WT, wild-type mice. n=3-7 animals per group.

### Visual responses are significantly reduced in *Rbpr2^-/-^* mice under vitamin A deficiency

We next recorded the electrical responses to light generated by rod photoreceptors, in dark-adapted mice eyes of *Rbpr2^-/-^* and WT mice, fed either a VAS or VAD diet, and the responses generated by cones in light-adapted eyes, by electroretinography (ERG) ranging from 0.01 to 100 cd.s/m^2^. The ERG analysis showed that scotopic *a*-wave and *b*- wave amplitudes were significantly decreased in 3-month old *Rbpr2^-/-^* mice on VAD diet, compared to *Rbpr2^-/-^* or WT mice on VAS diet (**Figures 5A-C**). This picture changed under long-term VAD diets. WT mice showed no changes in ERG amplitudes under either diets, however, *Rbpr2^-/-^* mice showed an even more severe decrease in *a*-wave and *b*- wave visual responses, compared to either *Rbpr2^-/-^* on VAS or WT mice on VAD and VAS diets at the 6-month analysis time-point (**Figures 5D-F**). Kinetic measurement of rod opsin recovery were also found to be slower in *Rbpr2^-/-^* mice on VAS or VAD diets, compared to WT mice on VAS diet, likely indicating decreased concentrations of the visual chromophore rhodopsin (**Figures 6A-6D**).

**Figure 5:**
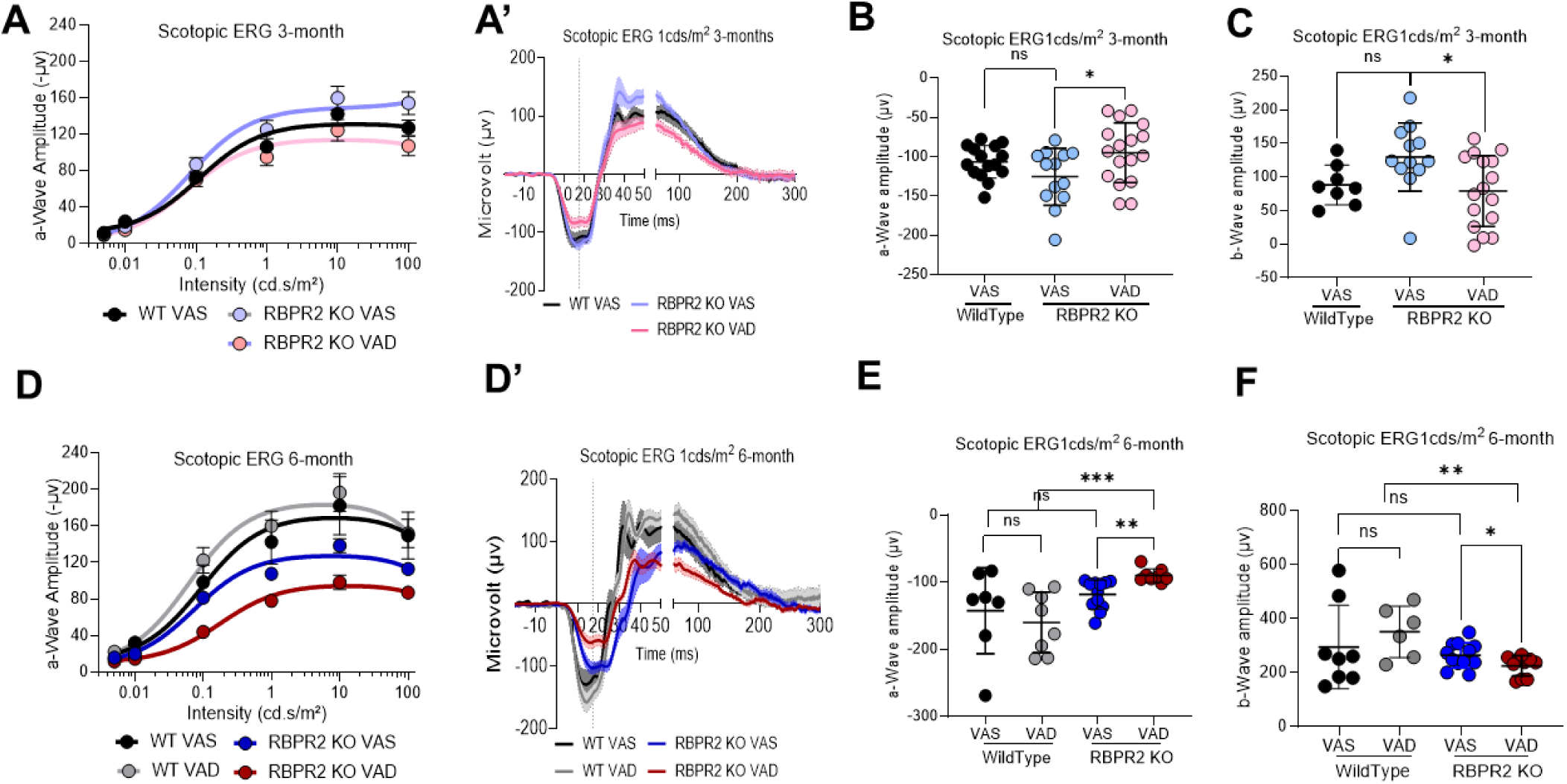
Rod photoreceptor cell functional analysis by Electroretinogram (ERG). Photopic ERG responses of WT and *Rbpr2^-/-^* mice at 3-months (**A-C**) and 6-months (**D-F**) of age fed either a VAS or VAD diet, showing the dark-adapted ERG responses of rod photoreceptor cell function by *a*-wave amplitudes (**B, E**) and inner neuronal bipolar cell function by *b*-wave amplitude (**C, F**). Values are presented as ±SD. Student *t*-test, *p<0.05; **p<0.005; ***p<0.001. VAS, vitamin A sufficient diet; VAD, vitamin A deficient diet; WT, wild-type mice. n=8-17 animals per group at 3-months of age; n=6-11 animals per group at 6-months of age.

**Figure 6:**
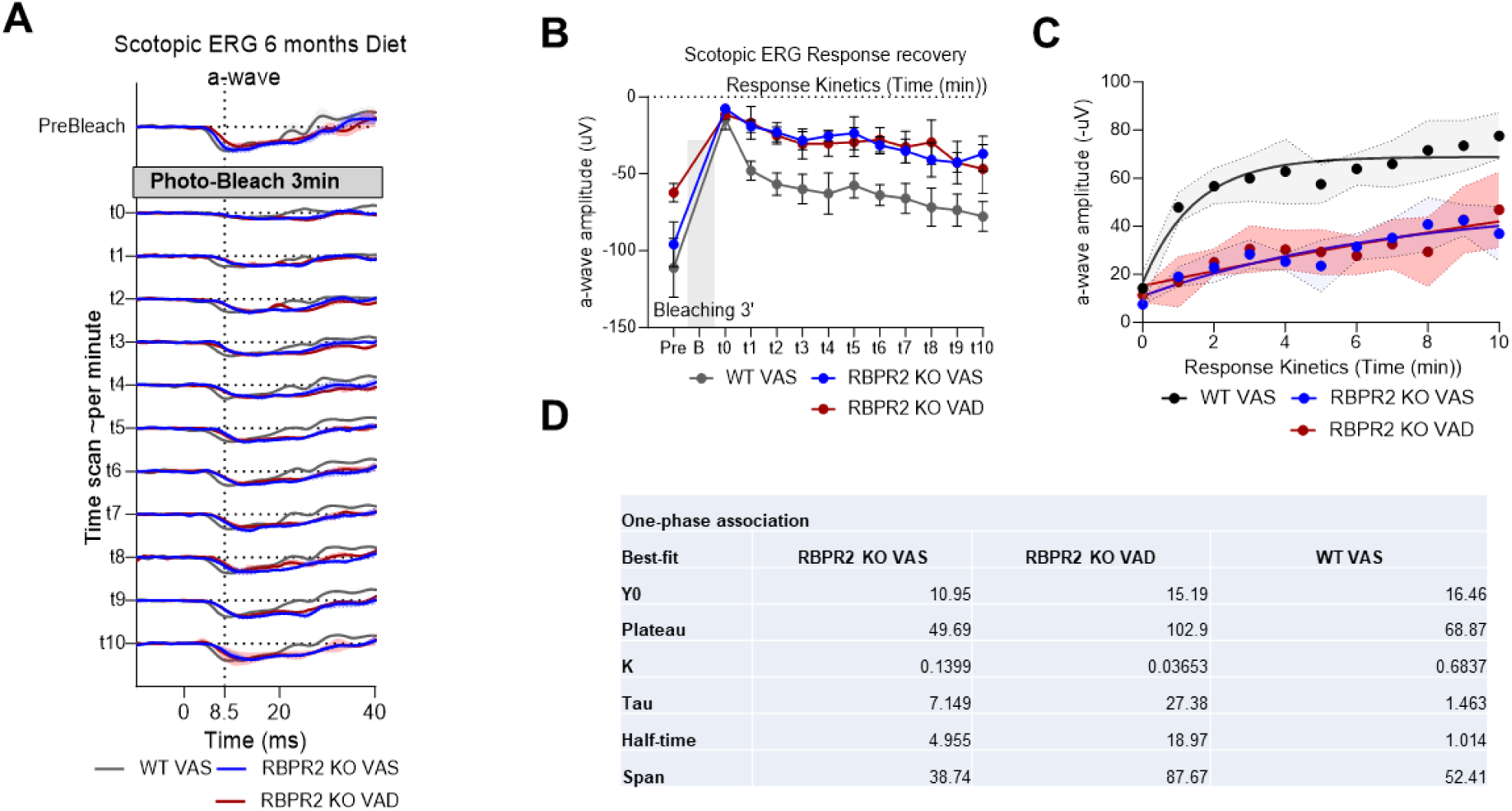
Rhodopsin physiological kinetics assessment by ERG photobleach recovery response. (**A**) Time series of ERG response in pre-bleaching, bleaching with full intensity blue stimulant light, and recovery response showing the kinetics curve of rod opsin in Wild Type and *Rbpr2^-/-^* mice fed with either a vitamin A sufficient (VAS) or vitamin A deficient (VAD) diet for six months. (**B-D**) One-phase association of a-wave amplitude showing the stability and half-life of rod opsin response recovery.

### Rod photoreceptors of *Rbpr2^−/−^* mice display significant levels of apoprotein opsin

Since *Rbpr2^−/−^* mice display reduced scotopic visual responses and decreased kinetics of rod opsin recovery, we hypothesized that this phenotype is caused by imbalances between chromophore and opsin concentrations in rod photoreceptors and likely to the presence of unliganded rod opsins or apoprotein opsins. It has been proposed previously that accumulation of apoprotein opsin/unliganded opsin can activate the phototransduction cascade even under dark conditions^13–15,18,32^. The constitutive activity of apoprotein opsin in photoreceptors is considered equivalent to background light and can result in a reduction in phototransduction gain^17,18,33^. We first analyzed the photoreceptor localization of rhodopsin in WT compared to *Rbpr2^-/-^* mice retinal sections by IHC at 6-months of age. In WT mice fed either a VAS or VAD diet, rhodopsin was properly localized to the photoreceptor outer segments (**Figure 7A**). Conversely, in *Rbpr2^-/-^* mice fed either a VAS or VAD diet significant (p<0.05) presence of mislocalized rod opsins were evident in the photoreceptor inner segments (**Figures 7A and 7B)**.

**Figure 7:**
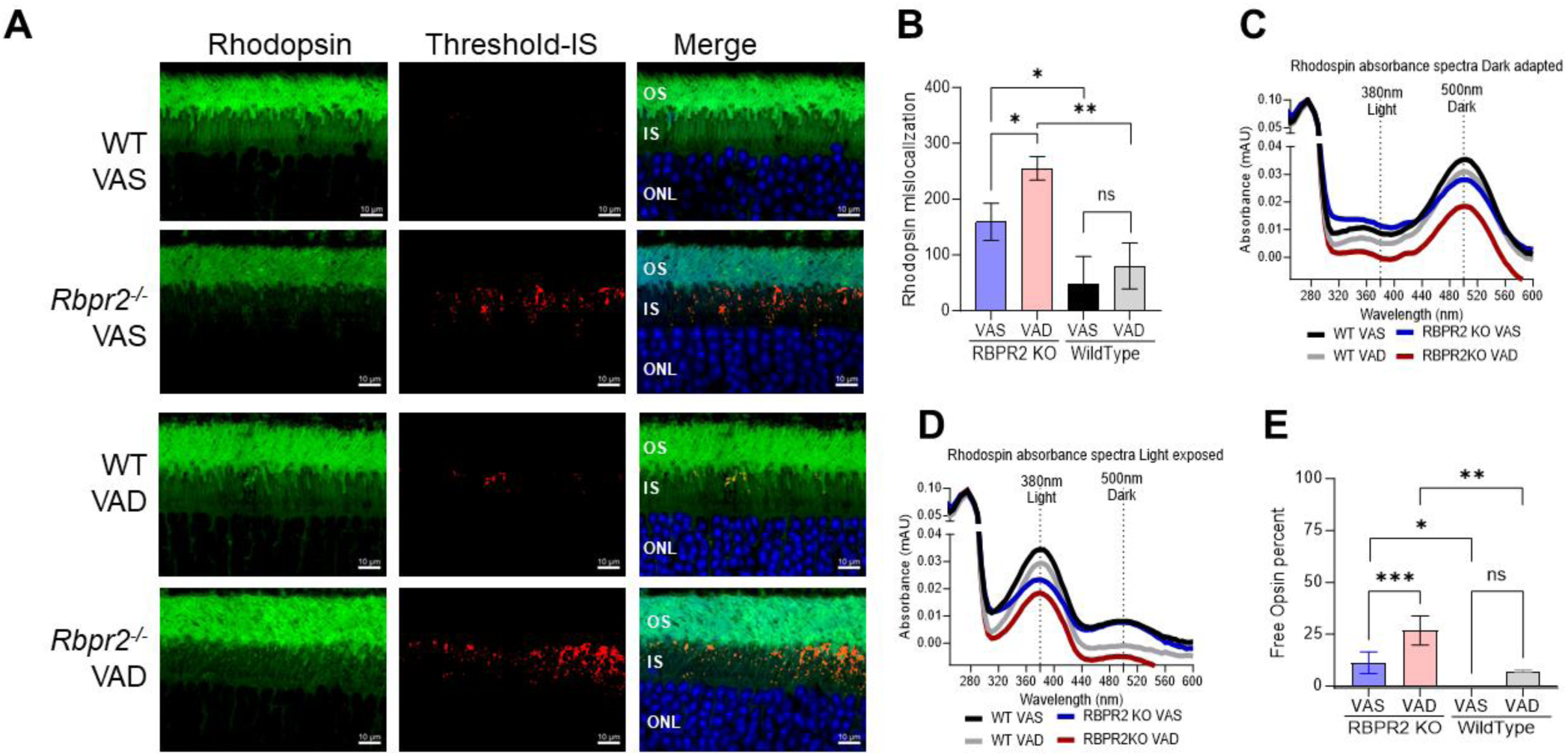
Presence of apoprotein opsin in rod photoreceptors of *Rbpr2^-/-^* mice. (**A**) IHC staining for Rhodopsin in photoreceptor OS in green and mislocalization in IS analyzed by the threshold in Red, Outer nuclear layer (ONL) stained with DAPI in Blue and merged images showing the localization of mislocalized Rhodopsin. (**B**) Threshold- based quantification of mislocalized Rhodopsin in the IS of retinas from WT and *Rbpr2^-/-^* mice fed different vitamin A diets. (**C**) UV-visible spectra of the immune-purified rhodopsin fractions from retinas of adult WT and *Rbpr2^-/-^* mice fed different vitamin A diets at the 6- month time point. Rhodopsin absorbance showing the peak absorbance at 500 nm for dark-adapted and 380 nm for 30-second high-intensity light-exposed samples (**D**). (**E**) Free opsin quantification shows the 11-*cis* retinal free apoprotein opsin percentage in the retinas of *Rbpr2^-/-^* mice compared to WT mice. VAS, vitamin A sufficient diets; VAD, vitamin A deficient diets. Values are presented as ±SD. Student *t*-test, *p<0.05; **p<0.005; ***p<0.001.

We next determined the levels of unliganded/ apoprotein opsin in *Rbpr2^−/−^* mice fed different vitamin A diets by performing UV-visible spectrophotometry with the isolated retinal protein fractions from WT and *Rbpr2^−/−^* mice using 1D4 antibody and compared the theoretical 280/500 nm ratio with the experimental ratio of absorbance^32^. We observed an ∼31% and ∼18% decrease in rhodopsin concentrations in dark-adapted photoreceptors of *Rbpr2^−/−^* mice fed either a VAD or VAS diet, compared to age-matched WT mice on VAS or VAD diets, respectively (**Figure 7C**). Similar results were observed in light-adapted photoreceptors, where lower Meta II rhodopsin concentrations were observed in *Rbpr2^−/−^* mice, compared to WT mice on either diets (**Figure 7D**). Quantification of unliganded opsin in dark-adapted retinas showed that *Rbpr2^−/−^* mice had significant amounts of apoprotein opsin, compared to WT mice on either diets (p<0.05, **Figure 7E**).

### Cone visual responses are significantly reduced in *Rbpr2-/-* mice on long-term vitamin A deficient diet

We next determined the ERG responses in light-adapted *Rbpr2^-/-^* mice fed either the VAS or VAD diets and using different light color sources (green, red, white, and UV/blue). Under photopic light conditions, ERG responses of *Rbpr2^-/-^* mice under green, white, and UV color sources were significantly diminished at the 3-month time point, while red light source ERG responses were not changed, when compared to WT mice on VAS diets (**Figures 8A-D**). Light-adapted ERG responses for green and white light intensity improved in *Rbpr2^-/-^* mice under VAS diets at the 6-month time point, but remained significantly diminished under blue/ UV-light exposure (**Figures 8A’-D’**). Kinetic measurement of cone opsin recovery under photopic blue light were slower in *Rbpr2^-/-^* mice on VAS or VAD diets, compared to WT mice on VAS diet (**Figure 9**). IHC for Red-green opsins (Opn1mw) in photoreceptors, showed that in WT mice fed either a VAS or VAD diet, cone opsins were properly localized to the photoreceptor outer segments (**Figure 8E**). Conversely, in *Rbpr2^-/-^* mice fed either a VAS or VAD diet significant (p<0.05) amounts of mislocalized cone opsins were evident in the photoreceptor inner segments (**Figure 8F)**.

**Figure 8:**
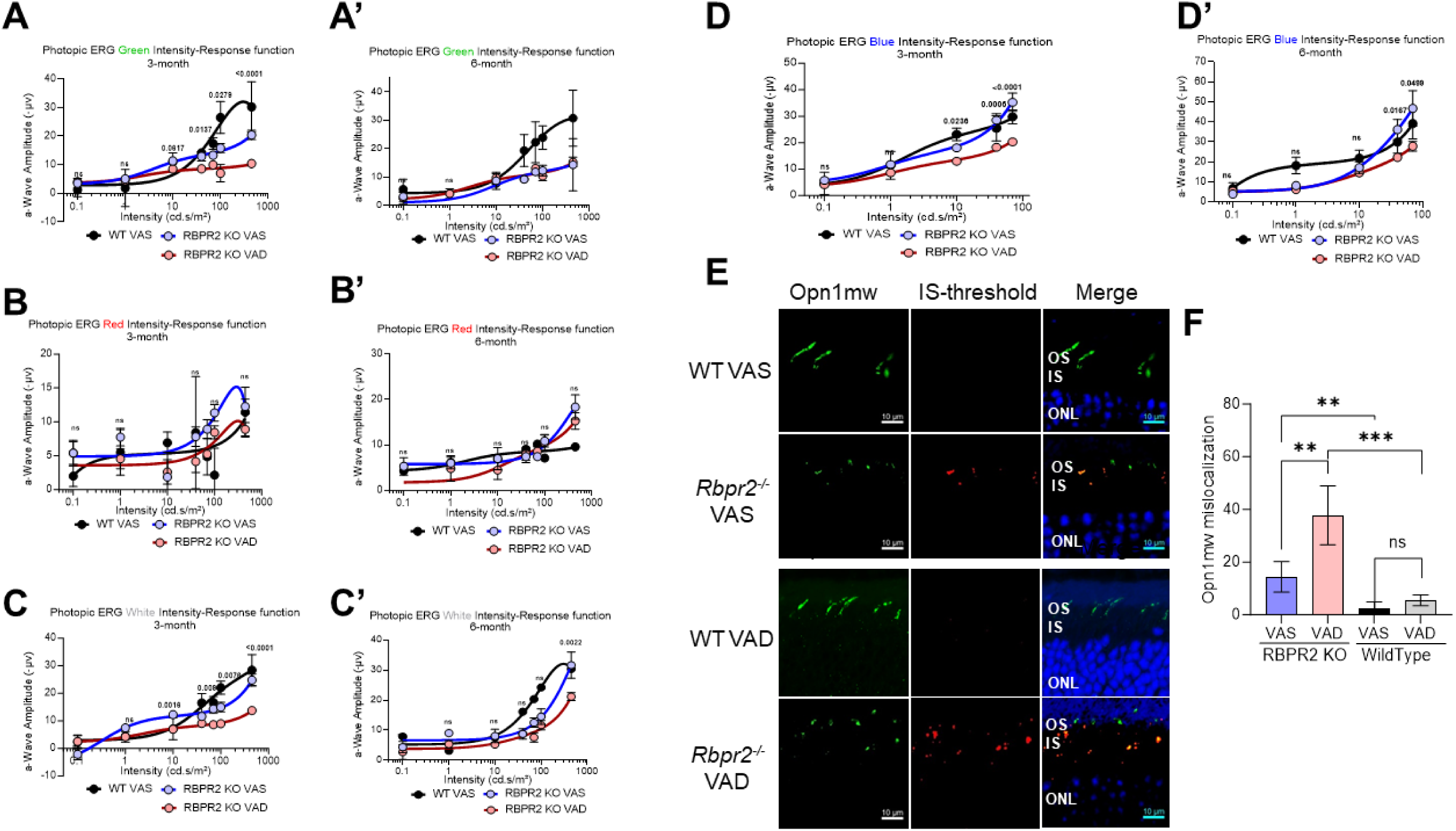
Cone Photoreceptor functional analysis by Electroretinogram. Photopic ERG of WT and *Rbpr2^-/-^* mice fed with either a VAS or VAD diet and stimulated with Green, Red, White, or Blue wavelength light intensities series showing the response curves, at 3-months of age (**A-D**) or 6-months of age (**A’-D’**). IHC staining for cone opsin (Opn1mw) in photoreceptor OS in green and mislocalization in IS was analyzed by the threshold in Red, Outer nuclear layer (ONL) stained with DAPI in Blue and merged images showing the localization of mislocalized cone opsins (**E**). Threshold-based quantification of mislocalized cone opsin in the IS of retinas from WT and *Rbpr2^-/-^* mice fed different vitamin A diets (**F**).

**Figure 9:**
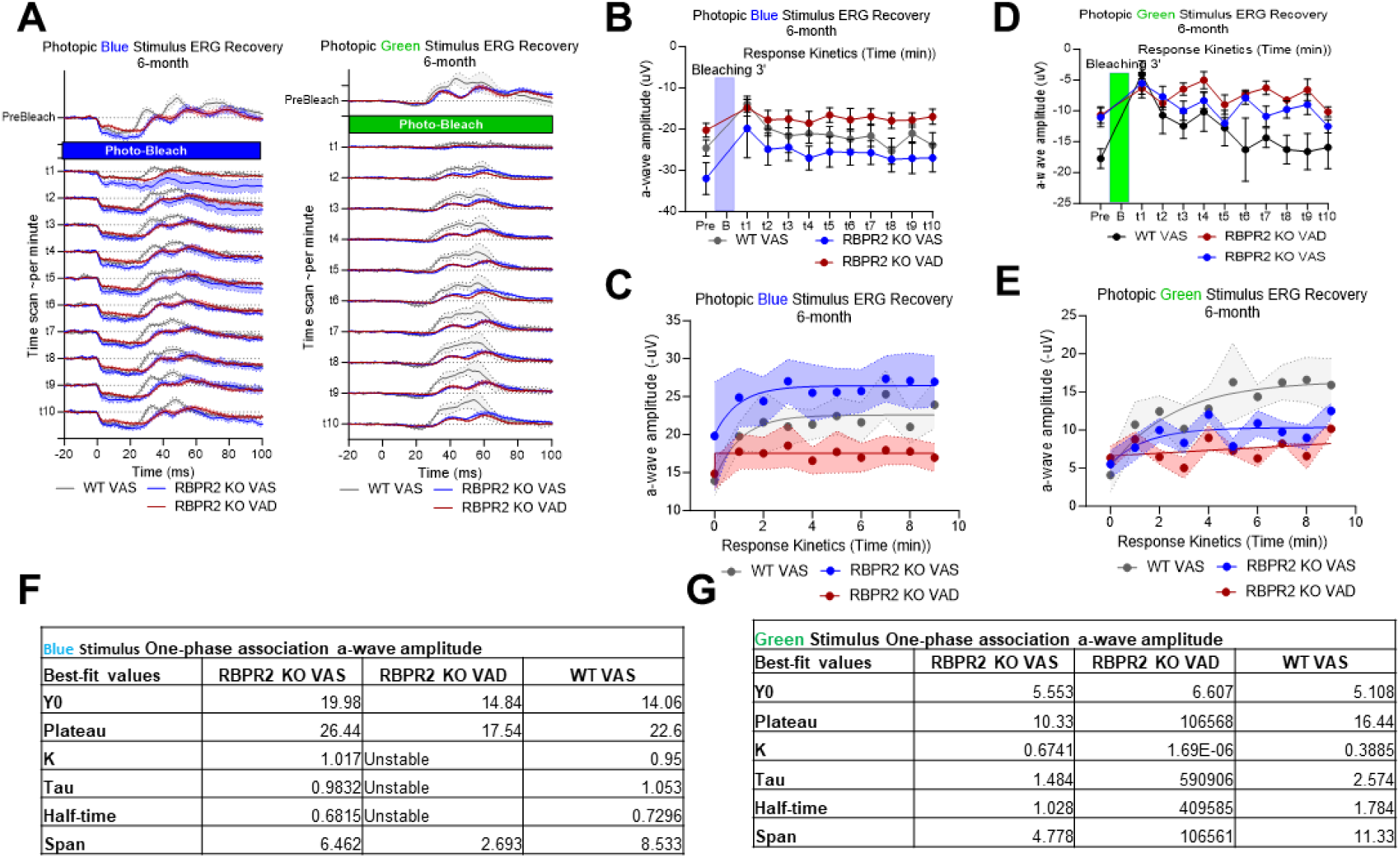
Short wavelength cone opsin Opn1sw physiological kinetics assessment by ERG photobleach recovery response. (**A-E**) time series of ERG response in pre-bleaching, bleaching with full intensity blue and green stimulant light and recovery response showing the kinetics curve in WT and *Rbpr2^-/-^* mice fed with either a VAS or VAD diet at 6-months of age. (**F, G**) One-phase association of *a*-wave amplitude and *b*-wave amplitude showing the stability and half-life of cone opsin response recovery.

## DISCUSSION

Dietary vitamin A (all-*trans* retinol/ROL) obtained from plant and animal sources is known to play an important role in metabolism, cell growth, immunity, reproduction, and visual function in humans^39–41^. Vitamin A deficiency (VAD) is a serious health issue, which is correlative with higher rates of mortality, childhood obesity, and nutritional blindness, especially among children in poorer countries around the world^39,40,42^. Vitamin A constitutes a group of biochemical compounds, including retinol, retinaldehyde, retinoic acid, and beta-carotene^39–41^. Most pertinent for visual function, 11-*cis* retinaldehyde (retinal) combines with the GPCR protein opsin in the photoreceptor outer segments to generate rhodopsin^2,4^. Prolonged VAD in the eye leads to impaired night vision due to deficient rhodopsin formation and can cause photoreceptor cell death and blindness^4^. Thus, an understanding of mechanisms that facilitate and regulate the uptake, transport, and long-term storage of ROL for systemic and ocular retinoid homeostasis is significant to the design of strategies aimed at attenuating retinal degenerative diseases associated with ocular ROL deficiency^6–18^.

In this study, we investigated the systemic and ocular consequences resulting from loss of the second RBP4-vitamin A transporter, RBPR2, on a longitudinal timescale. Previously, we have established a global *Rbpr2*-knockout (*Rbpr2^-/-^*) mice and have demonstrated that these mice are susceptible to visual deficiencies^19^. Here, we sought to expand upon that study in several critical ways. First, through a modified normal phase HPLC method, we are able to resolve retinaldehyde and retinol isomers, rather than just resolve total retinoid content. This is especially important for retinoid analysis in ocular tissues and allows us to directly detect and quantify critical retinoid isomers such as 11- *cis* retinal, the visual chromophore responsible for activation of the phototransduction cascade. Second, we have performed this analysis on multiple systemic tissues across multiple organs systems, rather than just in ocular tissue. Given that RBPR2 is hypothesized to regulate systemic vitamin A homeostasis, we aimed to examine the changes to retinoid content on a systemic level in these *Rbpr2^-/-^* mice. Third, to investigate the underlying causes for reduced electroretinogram responses in *Rbpr2^-/-^* mice, we utilized UV-Vis spectrophotometry to determine whether the ocular levels of vitamin A affect the stoichiometric balance between GPCR protein opsin and the visual chromophore, 11-*cis* retinal, in the photoreceptors.

All-*trans* retinol bound to RBP4 (RBP4-ROL) is the fundamental transport form of vitamin A found within the circulation, and its distribution must be tightly regulated in the support of multiple body functions including visual function. RBPR2 is the analogous RBP4-vitamin A receptor to ocular STRA6 and is expressed in major peripheral organs including the liver, the main storage organ for dietary vitamin A. Prior to its discovery and characterization in 2013, various groups of researchers have hypothesized about the existence of a mechanism that allows for the liver to intake circulatory RBP4-ROL. Now that the existence of RBPR2 is known and its molecular functions are characterized, investigations towards the understanding of its physiological functions has since been conducted. Given its ability to uptake circulatory RBP4-ROL, we hypothesize that RBPR2 could be responsible for systemic retinoid homeostasis, and that disruption of RBPR2 might result in altered retinoid levels in peripheral organs, including the eye.

To investigate if loss of RBPR2 affected retinoid concentrations on a systemic level, we performed HPLC analysis on various systemic organs, including the liver, on wildtype and *Rbpr2^-/-^* mice. Furthermore, we have performed this age-matched analysis at both the 3-month and 6-month timepoints. In all analyzed tissues, all-*trans* retinol (ROL) was the predominant (if not only) retinoid found in all analyzed systemic tissue, with the exception of retinyl esters in liver and retinaldehydes in ocular tissue. This is congruent with ROL being the predominant transport form of dietary vitamin A in mammalian organisms. Hence, quantification of ROL will serve as viable metric for determination of retinoid levels in these tissues. To provide a more intuitive representation of the normalized ROL quantification data, a heatmap plot of ROL levels across analyzed tissues, genotypes, and time points was generated. From this heatmap plot, a clear pattern emerges. Wild-type (WT) mice at both 3-month and 6-month timepoints are able to maintain ROL levels, for both VAS and VAD diets. However, while *Rbpr2^-/-^* mice at the 3-month timepoint for both VAS and VAD diets are able to maintain similar levels of ROL when compared to WT mice, the ROL quantity in *Rbpr2^-/-^* mice fed a VAD diet at the 6- month significantly decreases (Figure 3). Retinoid metabolism, like many other biochemical pathways, contains redundant pathways. In particular, circulatory retinyl esters within chylomicrons originating from the VAS diet provides the most likely explanation for maintenance of ROL levels comparable to WT in these *Rbpr2^-/-^* mice at both 3-month and 6-month time points, since this pathway bypasses the loss of *Rbpr2*\*\*\*. Similar, observations were obtained in *Stra6^-/-^* mice fed a high vitamin A diet^29,32^. However, for the *Rbpr2^-/-^* mice fed a VAD diet, this supplementary pathway does not exist. Once residual retinoid stores from gestation run out at the 6-month timepoint, these *Rbpr2^-/-^* mice fed a VAD diet exhibit greatly decreased ROL levels.

We next examined the retinoid supply and consumption axis in the support of visual function, by examining retinoid quantities in the liver and eye. While the liver was found to contain ROL like all other systemic tissues, the liver additionally contains considerable levels of retinyl palmitate, congruent with its role as the main storage organ of dietary vitamin A, with storage of retinoids predominantly in the form of retinyl esters within hepatic stellate cells. At both the 3-month and 6-month timepoints, hepatic ROL levels were found to be significantly lower for *Rbpr2^-/-^* mice on both VAS and VAD diets (Figures 2B and 4B). However, *Rbpr2^-/-^* mice fed the VAS diet generally exhibited comparable total hepatic retinoid levels when compared to WT mice fed a VAS diet at both the 3-month and 6-month time points, while *Rbpr2^-/-^* mice fed VAD diets exhibited significantly decreased hepatic total retinoid levels (Figures 2C and 4C). Under VAS conditions, both WT and *Rbpr2^-/-^* mice are continually converting retinyl ester stores into ROL for distribution into the bloodstream, thus exhibiting lower ROL levels but still displaying comparable total retinoid levels. In VAD conditions *Rbpr2^-/-^* mice are less capable in coping with vitamin A restriction, thus displaying both lower total retinoid and ROL levels in liver analysis.

This overall pattern of depressed levels of retinoids for *Rbpr2^-/-^* mice under VAD conditions in both systemic and hepatic tissue is thus reflected in ocular tissue, where the total retinoid content, ROL content, and 11-*cis* retinal content were found to be significantly lower (Figures 2F, 2G, and Figures 4F-4I). More critically, these depressed retinoid levels coexist with changes on the phenotypic level, with *Rbpr2^-/-^* mice under VAD conditions exhibiting mislocalized opsins within photoreceptor inner segments (Figures 7A-B), increased ratios of unliganded apoprotein opsin (Figures 7C-7E), and decreased rod and cone responses as measured with scotopic and photopic ERGs respectively (Figures 5, 6, 8, and 9).

In the past, investigations into mice exhibiting disruptions in the generation of 11- *cis* retinal has displayed phenotypes such as elevated levels of apoprotein opsins and subsequent retinal degeneration, such as in mice with disrupted *Stra6* and *Rpe65*^14,15,17,32^. In particular, *Rpe65^-/-^* mice, a mice model for the retinal degenerative disease Leber Congenital Amaurosis, has been shown to exhibit an elevated concentration of apoprotein opsin. RPE65 is an isomerohydrolase responsible for the conversion of retinyl esters to 11-*cis* retinol within the visual cycle, which is in turn necessary for the generation of 11-*cis* retinal chromophore in photoreceptors. The authors of that study have attributed the cause of retinal degeneration in *Rpe65^-/-^* mice to elevated levels of apoprotein opsin, which constitutively stimulates the phototransduction cascade though stimulation of transducin signaling, where disruption of transducin signaling partially rescues the retinal degenerative phenotype^15^. In a study investigating the phenotypic effects of *Stra6^-/-^* mice, which disrupts not only the intake of ROL from circulatory RBP4-ROL into the RPE, but also the subsequent generation of the 11-*cis* retinal chromophore, these mice exhibited elevated concentrations of apoprotein opsin and retinal degeneration much like *Rpe65^-/-^* mice, but additionally also mislocalization of rod and cone opsins within the photoreceptors^32^. These observations in *Stra6^-/-^* and *Rpe65^-/-^* mice were also reflected in *Rbpr2^-/-^* mice, where significant rhodopsin mislocalization was observed in these mice fed VAS or VAD diets, but not in WT mice even on the VAD diet (Figures 7A, 7B). Moreover, studies examining the phenotypic effects of *Stra6^-/-^* mice additionally studied the effects of applying pharmacological doses of vitamin A as a means rescuing the ocular phenotypes in these mice. These studies show that interventions with high doses of ROL increase the total retinoid content found within ocular tissues, despite lacking a vitamin A membrane receptor to access circulatory RBP4-ROL due to its lack of functional STRA6. This is a result that was also reflected in *Rbpr2^-/-^* mice, where a VAS diet is able to supplement systemic retinol levels despite the disruption of vitamin A homeostasis through disruption of RBPR2^29,32^. As mentioned above, the retinoid delivery system in mammalian organisms exhibits plasticity in redundancy, where chylomicron transport of retinyl esters originating from the diet can act as a possible alternate pathway for delivering retinoids to systemic tissues^19^.

Given that RBPR2 is hypothesized to function as a systemic regulator of vitamin A homeostasis and that *Rbpr2^-/-^* mice display similar ocular phenotypes as other mouse models with disrupted chromophore generation, including apoprotein opsin accumulation, decreased scotopic and photopic responses, and opsin mislocalization, our study indicates that RBPR2 is an important facilitator of visual function through its mechanistic role as a systemic regulator of vitamin A homeostasis.

## AUTHOR CONTRIBUTIONS

Conceptualization, G.P.L.; writing-original draft preparation, G.P.L., R.R., and M.L., performed experiments, R.R., A.L., S.M., and M.L., manuscript writing, review, and editing, G.P.L., R.R., S.M., M.L.; supervision, G.P.L.; project administration, G.P.L.; funding acquisition, G.P.L. All authors have read and agreed to the published version of the manuscript.

## AKNOWLEDGEMENTS

We thank Dr. Ahmed Sadah M.D., for his assistance with mice colony maintenance and genotyping, and Dr. Beata Jastrzebska, Ph.D. (Department of Pharmacology, Case Western Reserve University, OH) for her advice with the rhodopsin absorbance protocol.

## Abbreviations

ROL: all-*trans* retinol
11-cis RAL: 11-cis retinaldehyde
RBPR2: Retinol binding protein 4 receptor 2
LRAT: Lecithin:ROL acyltransferase
*at*RA: all-*trans* retinoic acid
STRA6: stimulated by retinoic acid protein 6
RBP4: Retinol binding protein 4
Stra6l: Stimulated by retinoic acid protein 6 like
HPLC: High Performance Liquid Chromatography

---

**Supplementary Figure S1:**
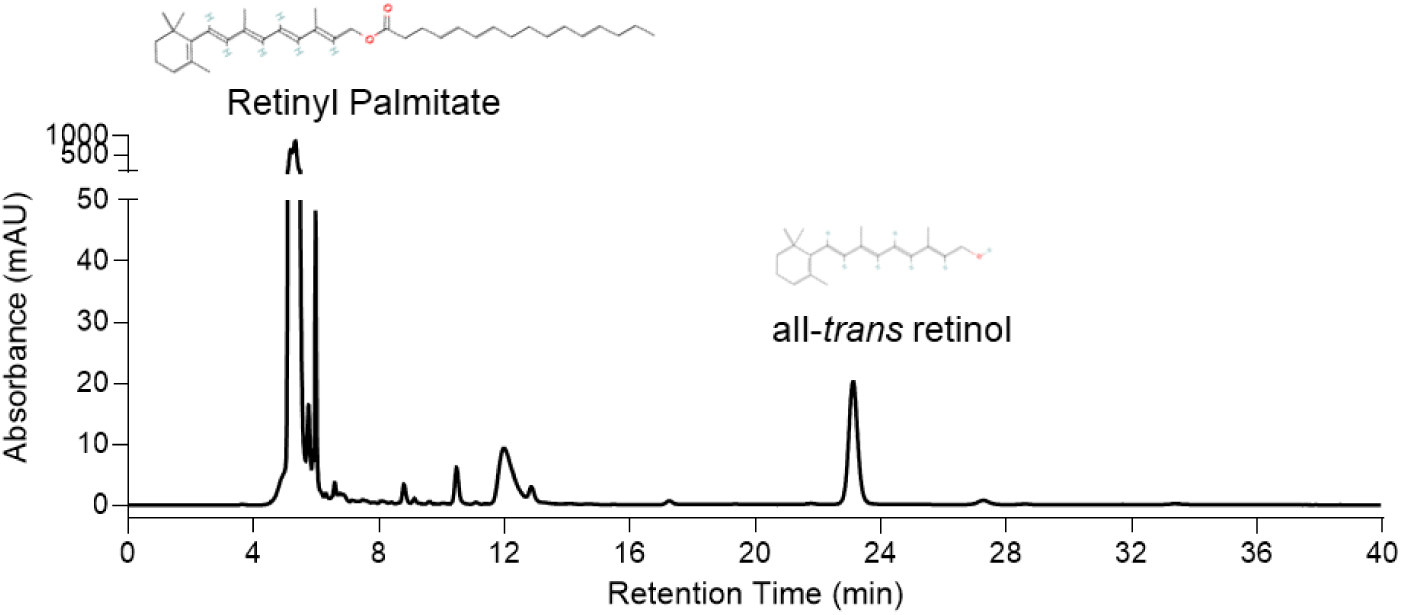
Representative HPLC chromatogram of retinoids from Wild- type mice liver.

**Supplementary Figure S2:**
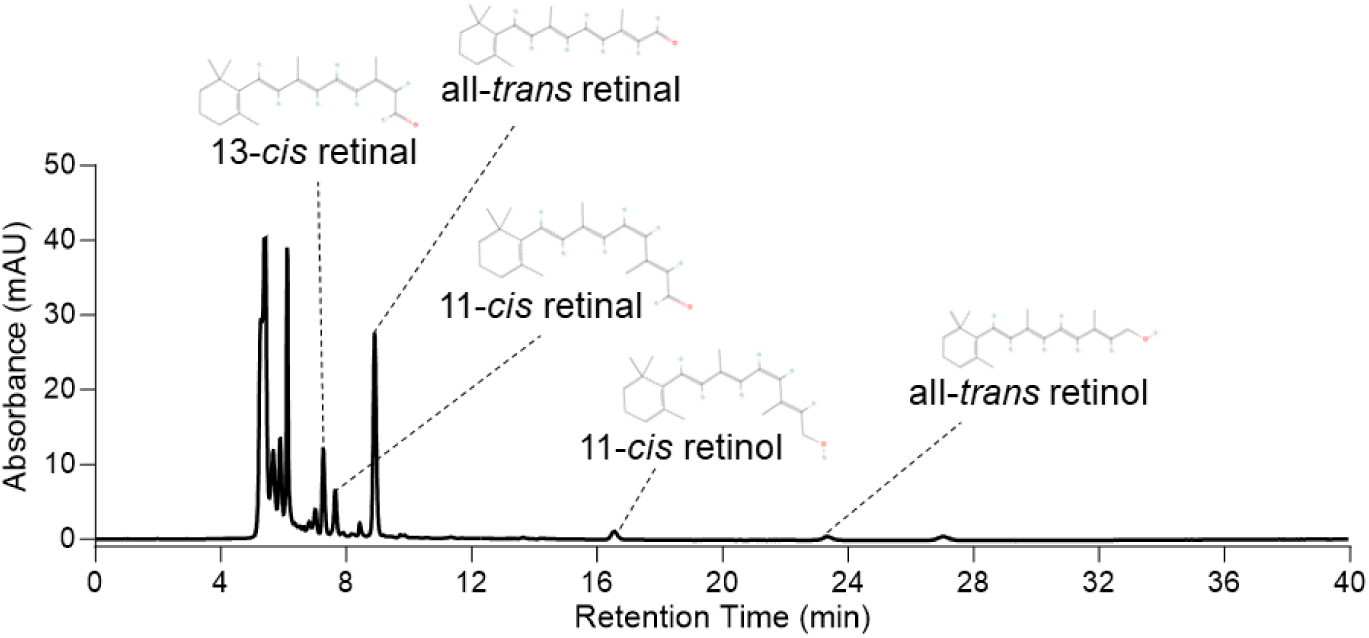
Representative HPLC chromatogram of retinoids from wild- type mice eyes.

**Supplementary Figure S3:**
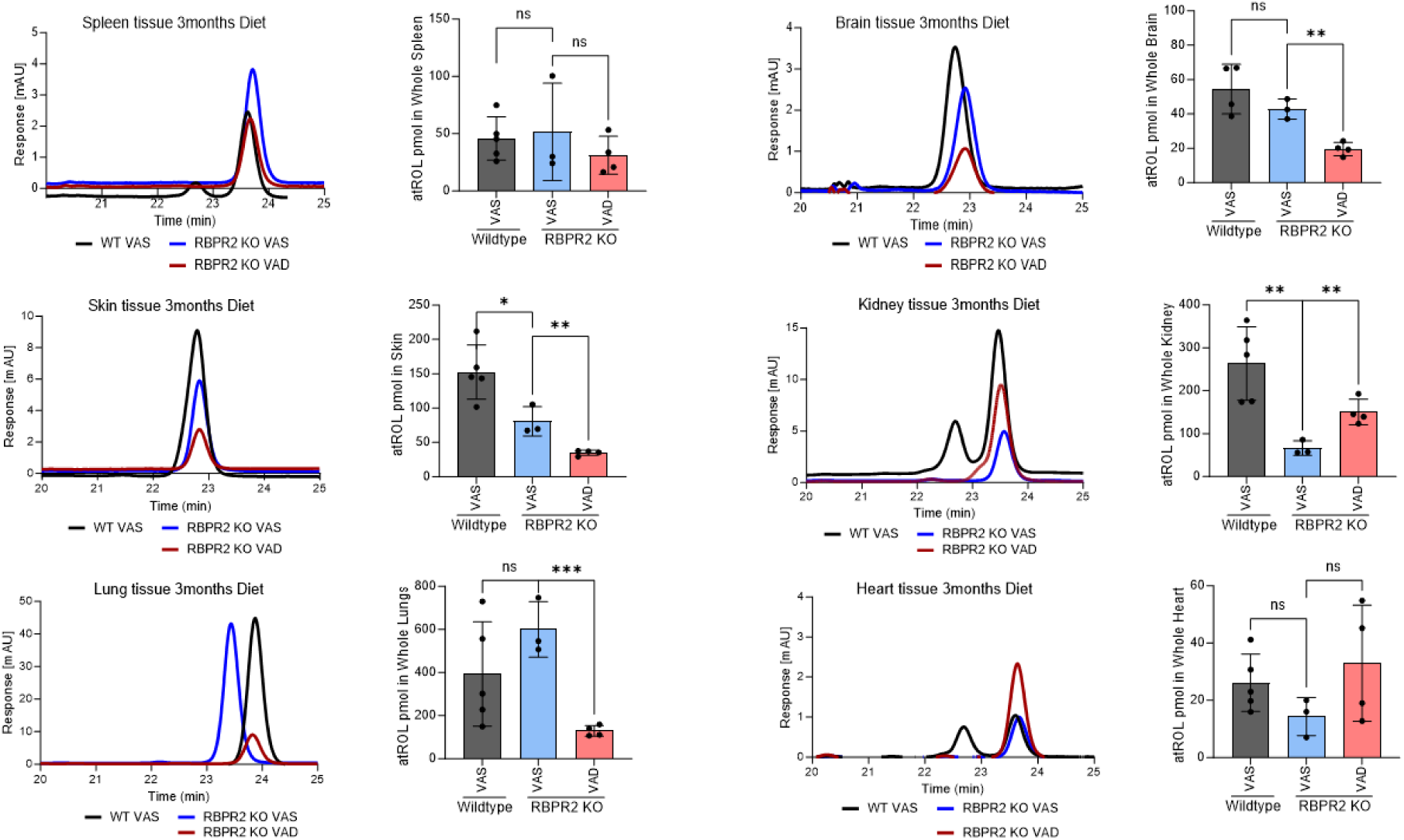
HPLC analysis and quantification of all-*trans* retinol at 3-months of age in various tissues. WT and *Rbpr2^-/-^* mice on different vitamin A diet showing the comparative box plots of all-*trans* retinol (atROL) in various non-ocular tissues among the genotypes and dietary conditions. Values are presented as ±SD. Student *t*-test, *p<0.05; **p<0.005; ***p<0.001; n.s., not significant.

**Supplementary Figure S4:**
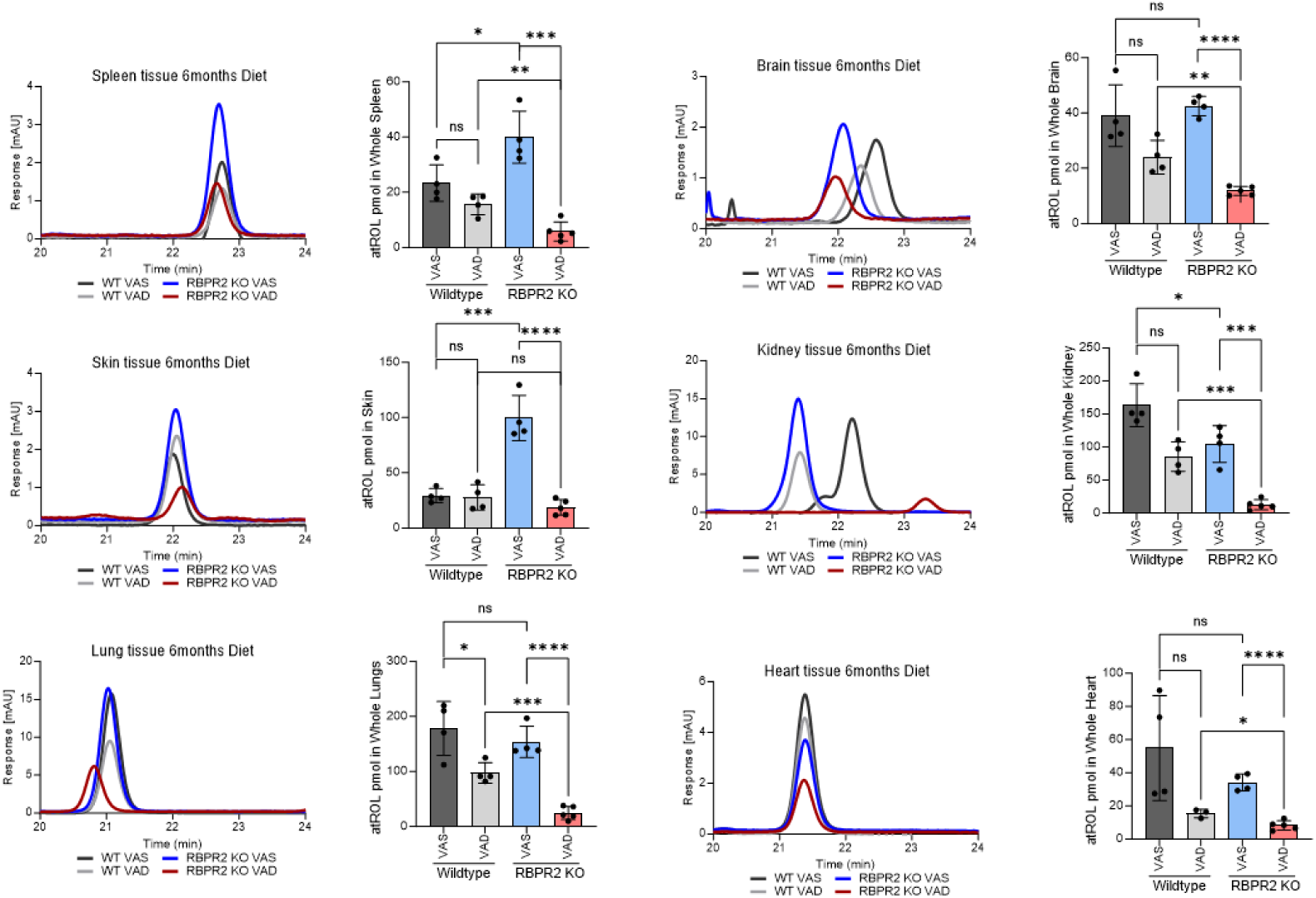
HPLC analysis and quantification of all-*trans* retinol at 6-months of age in various tissues. WT and *Rbpr2^-/-^* mice on different vitamin A diets showing the comparative box plots of all-*trans* retinol (atROL) levels in various non-ocular tissues among the genotypes and dietary conditions. Values are presented as ±SD. Student *t*-test, *p<0.05; **p<0.005; ***p<0.001; ****p<0.0001.; n.s., not significant.

