## SUPPLEMENTARY MATERIAL for "Loss of the systemic vitamin A transporter RBPR2 affects the quantitative balance between chromophore and opsins in visual pigment synthesis"

### SUPPLEMENTARY FIGURES

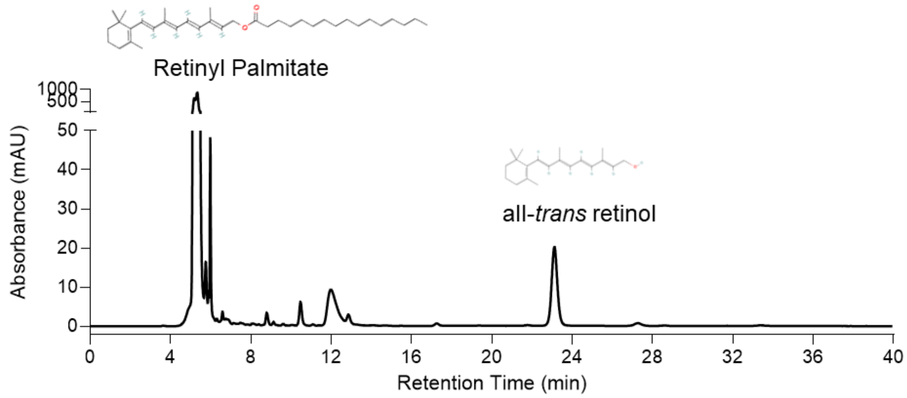


**Supplementary Figure S1:** Representative HPLC chromatogram of retinoids from Wild-type mice liver.

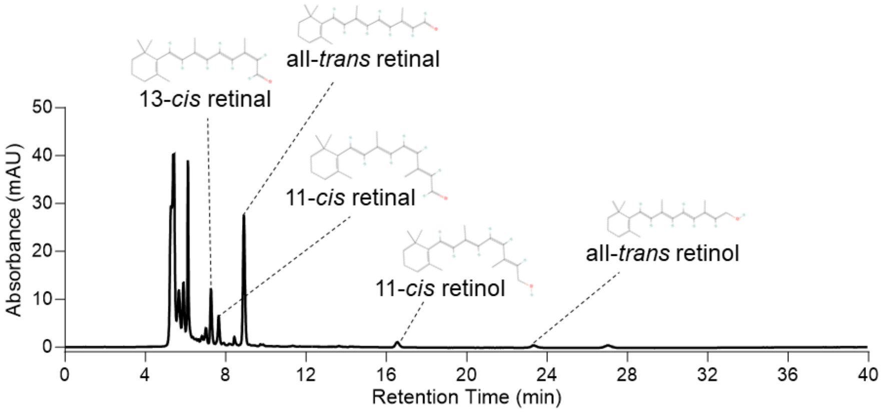


**Supplementary Figure S2:** Representative HPLC chromatogram of retinoids from wild-type mice eyes.

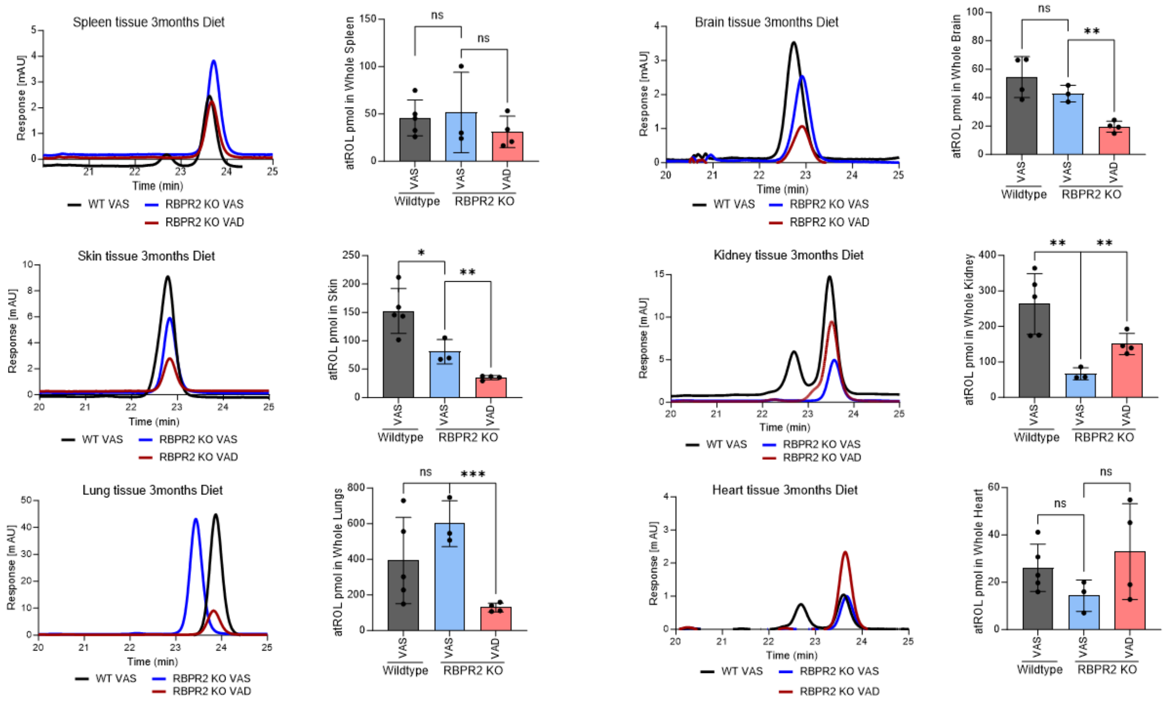


**Supplementary Figure S3: HPLC analysis and quantification of all-*trans* retinol at 3-months of age in various tissues.** WT and *Rbpr2*$^{-/-}$ mice on different vitamin A diet showing the comparative box plots of all-*trans* retinol (atROL) in various non-ocular tissues among the genotypes and dietary conditions. Values are presented as ±SD. Student *t*-test, *p<0.05; **p<0.005; ***p<0.001; n.s., not significant.

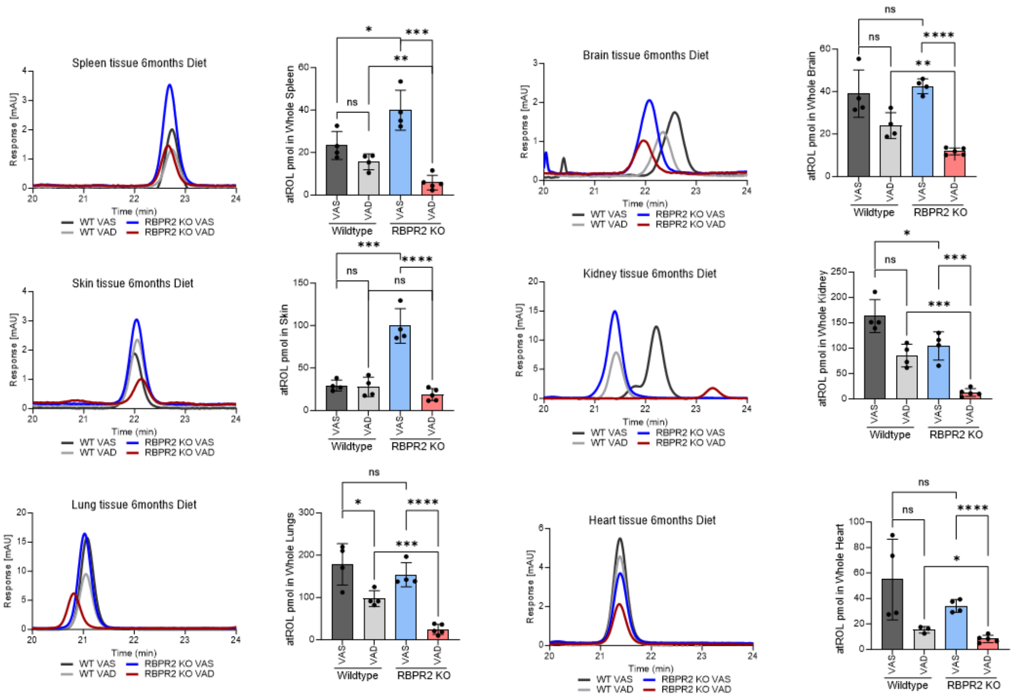


**Supplementary Figure S4: HPLC analysis and quantification of all-*trans* retinol at 6-months of age in various tissues.** WT and *Rbpr2*$^{-/-}$ mice on different vitamin A diets showing the comparative box plots of all-*trans* retinol (atROL) levels in various non-ocular tissues among the genotypes and dietary conditions. Values are presented as ±SD. Student *t*-test, *p<0.05; **p<0.005; ***p<0.001; ****p<0.0001.; n.s., not significant.
